# Prediction, Syntax and Semantic Grounding in the Brain and Large Language Models

**DOI:** 10.1101/2025.06.05.658007

**Authors:** Nikola Kölbl, Stefan Rampp, Martin Kaltenhäuser, Konstantin Tziridis, Andreas Maier, Thomas Kinfe, Ricardo Chavarriaga, Patrick Krauss, Achim Schilling

## Abstract

Language comprehension involves continuous prediction of upcoming words, with syntactic structure and semantic meaning intertwined in the human brain. To date, few studies have used combined magnetoencephalography (MEG) and electroencephalography (EEG) measurements to investigate how syntactic processing, predictive coding, and semantic grounding interact in real time. Here we present the first combined MEG-EEG investigation of syntactic processing and semantic grounding under naturalistic conditions. Twenty-nine healthy participants listened to a German audio book while their neural responses were recorded. Event-related fields and event-related potentials for four word classes - nouns, verbs, adjectives, and proper nouns - showed highly reproducible, characteristic spatio-temporal signatures, including significant pre-onset activity for nouns, suggesting enhanced predictability of this word class. Source-space analyses revealed pronounced activation in the pre- and post-central gyri for nouns, suggesting a deeper semantic grounding of nouns in e.g. sensory experiences than verbs. To further investigate predictive mechanisms, we analyzed the hidden representations of the large language model Llama. By comparing the transformer-based representations to neural responses, we explored the relationship between computational language models and human brain activity, offering new insights into syntactic and semantic prediction. These findings highlight the power of simultaneous MEG-EEG recordings in unraveling the predictive, syntactic, and semantic mechanisms that underlie the comprehension of natural language.

## 1 Introduction

Emerging from cutting-edge AI research, large language models (LLMs) such as GPT4o and Llama have revolutionized the way we interact with machines, surprising users with their human-like responses [San23; Dal21; Bro+20; Tou+23b]. These models rely almost entirely on predictive processes to generate language [Ewe+24; Qi+24], enabling them to approach - or even surpass - the Turing-Test with unprecedented proficiency [Pro24; Bor25].

The human brain, too, is not a passive receiver, but an active “prediction machine”, constantly anticipating upcoming words and events [Bar09; SK24; Sch+23a]. Language - perhaps the most central of all human capacities - underpins our culture, society, science, and collective progress [NK99; Kra+24a]. Understanding the neural mechanisms that generate such linguistic predictions can provide insights for casting light on the very foundations of human communication and cognition.

Despite significant progress in uncovering the neural basis of language, it remains largely elusive how the brain stores and processes grammar and grammatical structures [Bog96; Din+16] - do they emerge through predictive use, or are they innately pre-specified? Despite our increasingly deep understanding of these predictive processes, questions about how different grammatical frameworks are represented, stored, and ultimately learned continue to fuel a vigorous debate [Fri+17; GPS22].

Currently, two prominent schools of thought - Chomsky’s theory of universal grammar [Cho98; Cho14] and the usage-based cognitive linguistics approach [Gol95; Gol03; Tom05; Lan08] - offer competing explanations of how grammatical structures are acquired and represented in the brain. Chomsky’s theory of universal grammar posits that humans possess an innate, biologically determined language ability that enables rapid identification of word categories, such as nouns and verbs, and thus facilitates language acquisition in children [Cho98; Cho14; Yan+17]. In contrast, cognitive linguistics emphasizes a deep interplay between language structure and language use, suggesting that grammar emerges as a dynamic system shaped by contextual processing and mental representations [Gol95; Gol03; Tom05; Lan08].

Moreover, syntax directly shapes meaning, as, e.g., the position of a word in a sentence can change the semantic content of the whole sentence [GIM13] - a fact that is taken into account in LLMs by the positional encoding method (see e.g., [Tou+23b; Tou+23a]). Thus, if language indeed represents the fundamental ability required for the development of general intelligence, chain-of-thought reasoning and abstract cognition, and if grammar naturally emerges through language usage — thereby aligning brain mechanisms with patterns observed in deep neural networks — this raises the critical question: can LLMs trained solely on next-word prediction evolve into artificial general intelligence (AGI) [Fen+24]?

According to the “symbol grounding problem” [Har90; TF05], this is very unlikely, as the meaning of words these neural networks manipulate is not grounded in the real world. Therefore, to better understand how these mechanisms work in the human brain and how they could be adapted for artificial intelligence, syntax processing and semantic grounding should not be regarded in isolation [DD16].

Resolving these long-standing issues will likely require a two-pronged strategy: on the theoretical front, using cutting-edge machine learning algorithms as computational models of brain function [KD18; KSK24; Met+24; Imm+24; RSK24b; RSK24a; Sto+24; Sto+22; Sto+23a; Sto+23b; Kra+24a; Ger+23], and on the empirical side, conducting precise neuroscience measurements with high temporal resolution (e.g., using EEG, MEG techniques) to study the neural correlates of language processing [Sch+21a; Sch+24; Sch+23b; Koe+24; KSK23].

From the theoretical side, we have already started to tackle this problem in a previous study by training a small five-layered brain-constrained recurrent language model on next-word prediction [Sur+23]. By examining the hidden representations of this model after training, we observed the spontaneous emergence of distinct clusters of activation in the internal feature space corresponding to specific word types [Sur+23]. This suggests that even a relatively simple neural network trained only on next-word prediction can spontaneously internalize basic grammatical structures. Hence, it seems plausible that the human brain, with its approximately 100 billion neurons, could accomplish this feat through continuous language-based prediction alone [Len+12; Ger+20; Sur+23].

However, syntax is not equivalent to semantics, and a purely computational account remains incomplete without empirical validation from experimental neuroscience, underscoring the need for direct measurements of how the human brain encodes and processes grammatical structures. Furthermore, it is necessary to move from simple neural networks as described above to state-of-the-art cognitive models, which are already able to solve complex cognitive and reasoning tasks such as LLMs [KD18; YP24; Bub+23].

Therefore, in the present study, we performed combined MEG/EEG measurements on the one hand and compared the neuronal patterns found to the patterns observed in the Meta’s Llama 3.2 LLM [AIM24; Tou+23b; Tou+23a].

Up to now, much of our understanding of language processing in the brain has come from neuroimaging studies that rely on carefully controlled, often simplified linguistic stimuli - such as single words, isolated sentences, or short phrases [ZK18; MPH16]. Although these paradigms allow researchers to pinpoint specific cognitive processes underlying comprehension, they fail to capture the dynamic complexity of natural language and can be heavily influenced by experimental design [Ber17]. Consequently, findings from such controlled contexts do not always generalize to everyday language use [Ald19]. In response, neurolinguistic research has increasingly adopted more realistic, continuous speech stimuli, including excerpts from audio-books, to better reflect real-world language processing [Sch+21a; KSK23; Gar+22; Sch+23b; Sch+24]. Such more realistic, context-enriched stimuli offer a promising way to immerse participants in the complexities of naturalistic language processing [Sch+21a].

In the present study, we used a naturalistic continuous speech stimulus and performed combined EEG and MEG measurements to investigate neural responses to four different word classes - nouns, verbs, adjectives, and proper nouns. To this end, we presented the first 6.5 chapters of two stories from the German science fiction audio-book “Vakuum” by Phillip P. Peterson to 29 healthy, right-handed german native speakers, allowing us to capture brain responses in a fluid linguistic environment [Sch+21a; Koe+24]. Inspired by machine learning protocols that use training and test sets to promote reproducible model development, we divided participants into exploration and validation cohorts, an approach that allowed for the explorative discovery of neural patterns associated with different word classes and their subsequent verification in an independent sample, thereby increasing the reliability and generalizability of our findings [Sch+21a; Koe+24; Kra+24b].

By incorporating this strategy, we aimed to enhance the reproducibility of our findings — a core aim of the “replication movement” in cognitive neuroscience [MHH18; Gil+17]. Finally, we also compared the recorded brain activity with outputs from the large language model LLama 3.2, probing for potential parallels in human and machine-based language processing.

Our analyses of event-related potentials (ERPs) and fields (ERFs), as well as spatio-temporal patterns in source space, reveal distinct processing profiles for different word classes from the earliest neural time windows. In particular, we observed significant pre-onset activity for nouns, suggesting heightened readiness for this word category - an effect consistent with predictive coding.

Moreover, in a complementary analysis, we trained a linear probe neural network on the hidden states of Llama 3.2, revealing that nouns and adjectives are more easily predicted than verbs — an outcome that aligns with the pre-onset ERFs and ERPs signals observed in our data. We discuss our findings in the context of predictive coding frameworks in artificial neural networks, highlighting potential parallels between biological and computational approaches to language anticipation.

## 2 Material and Methods

### 2.1 Participants

We recorded the neural responses of 29 participants (15 females, mean age 22.8 years, range 18-28 years) while they listened to an audio book. All participants were healthy, right-handed individuals (mean laterality quotient 85.4 ± 12.6 [Old71]), German native speakers with normal hearing, and reported no history of neurological disorders or substance abuse. The study protocol was approved by the Ethics Committee of the University Hospital Erlangen (No: 22-361-2, PK).

### 2.2 Experimental Design

As a natural language stimulus, we used about 50 minutes of the science fiction audio book “Vakuum” by Phillip P. Peterson (published by Argon Hörbuch, narrated by Uve Treschner). The text consists of several story-lines, two of which were selected and alternated in approximately seven-minute segments across eight chapters. After each chapter, participants answered three single-choice questions about the content to ensure their continued attention. Simultaneous recordings of brain activity were obtained with a 248-channel magnetoencephalography (MEG) system (Magnes 3600WH, 4D-Neuroimaging) and a 64-channel electroencephalography (EEG) system (ANT Neuro), supplemented with electrooculogram (EOG) and electrocardiogram (ECG) measurements.. To limit electromagnetic artifacts affecting the MEG sensors, sound was delivered through air tubes from loudspeakers placed outside a magnetically shielded chamber (see [Sch+21a]). The sound level was individually adjusted for clarity and comfort. Throughout the measurement, participants lay still and focused on a fixation cross to minimize eye and muscle artifacts.

### 2.3 Data Preparation

To improve the signal quality of both EEG and MEG datasets, we applied a standard pre-processing pipeline [Fer+22] using the MNE software (version 1.8.0, [Gra+13]). First, we identified and interpolated faulty sensors and electrodes - those with flat (zero variance) or excessively noisy (high variance) amplitudes. Next, we applied a 1-20 Hz band-pass filter to constrain the analysis to a relevant frequency range and down-sampled the data to 200 Hz to decrease computational complexity. We then performed Independent Component Analysis (ICA) to remove artifacts. Specifically, in addition to discarding the first two independent components (ICs) with the highest variance, we eliminated any ICs, which correlated with the simultaneously recorded EOG or ECG channels, thereby mitigating eye movement and heartbeat artifacts[KSK23].

To align and segment the continuous EEG and MEG recordings, we applied forced alignment (*WebMAUS* software [Sch15]) to the audio files and corresponding transcripts, extracting precise onset times for each word in the audio book. Simultaneously with the audio playback, we routed the audio signal to a dedicated stimulus channel in the MEG acquisition device (trigger channel). Using this channel and the word-onset information, we then segmented the MEG and EEG data into epochs from 1.0 s before to 2.0 s after each word onset (with baseline correction applied from -1.0 s to 0.0 s). Since our primary interest is in syntactic processing, we used the natural language processing software *spaCy* (model: “de_core_news_sm”) to assign part-of-speech (POS) tags to each word [Hon+20]. From these tags, we focused on four categories - nouns, verbs, adjectives, and proper nouns. Figure 1 illustrates both the distribution of word lengths across these categories and their respective frequencies in the audio book.

**Figure 1:**
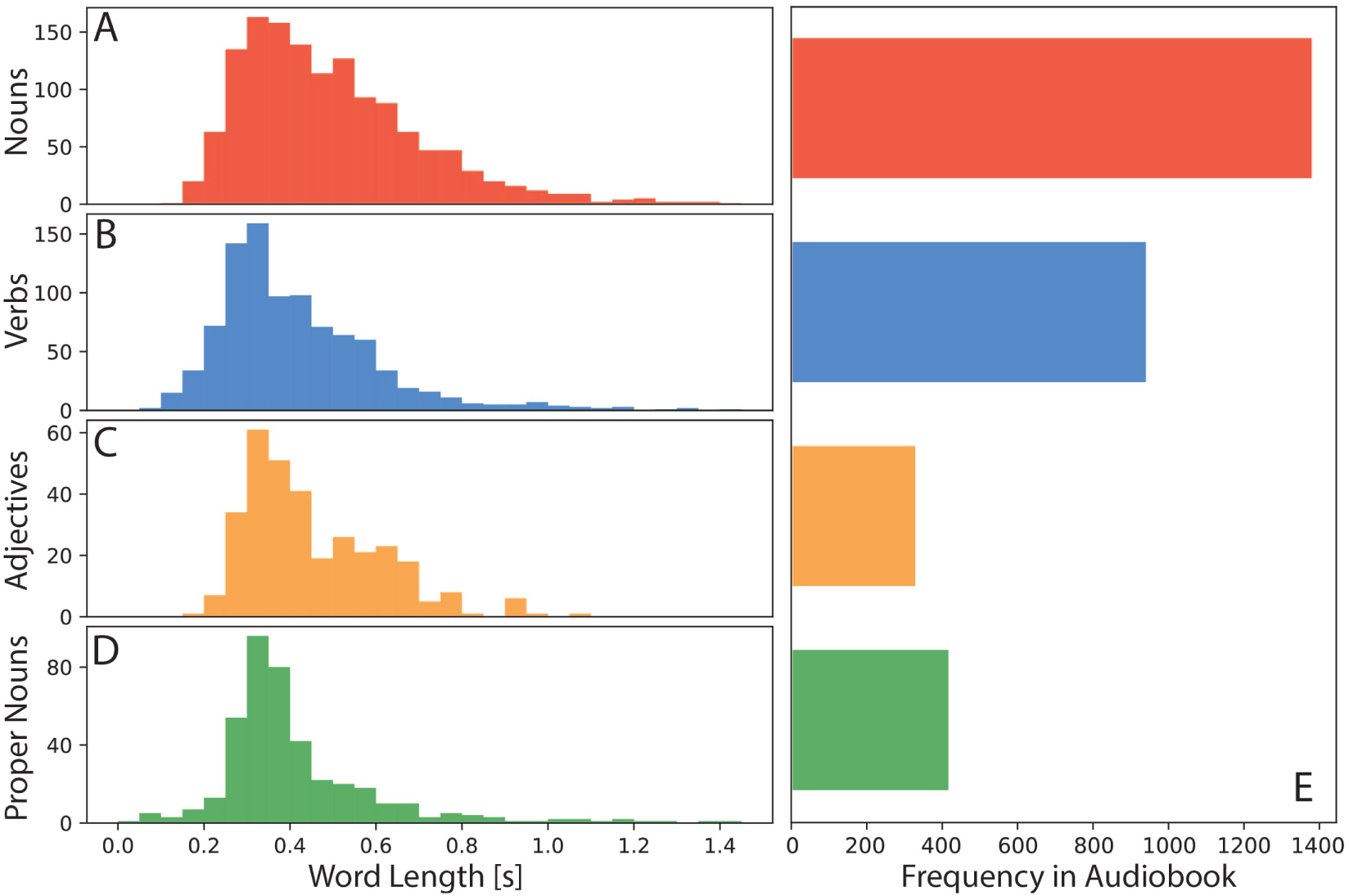
Distribution of word lengths for each word class. (A): Nouns, µ: 0.498s, *σ*: 0.220s. (B): Verbs, µ: 0.420s, *σ*: 0.198s. (C): Adjectives, µ: 0.457s, *σ*: 0.177s. (D): Proper Nouns, µ: 0.427s, *σ*: 0.228s. (E): Word-type frequency histogram in audio book. Nouns (red): 1376, Verbs (blue): 936, Adj (orange): 325, Proper Nouns (green): 413.

We further refined the word-type-specific event-related fields (ERFs) and event-related potentials (ERPs) by filtering them into the 1-4 Hz range using a one-pass, zero-phase finite impulse response (FIR) filter. Before analyzing the data, we divided the participants into two independent cohorts for exploration (subjects 1-19) and validation (subjects 20-29) to identify word-type-specific patterns in the grand-average-ERFs and ERPs that would prove consistency across the two groups. The subject numbers reflect the chronological order in which recordings were done (exploration set: 13 females, mean age 22.1 years, range 18-27 years; validation set: 2 females, mean age 24.0, range 19-28).

### 2.4 Source Reconstruction

For source reconstruction and subsequent data analysis, we used the open-access Brainstorm software [Tad+11], using the standard ICBM 152 anatomy from the MNI database as our head model [Fon+09].

Prior to each EEG-MEG measurement, we used a Polhemus digitizer to record fiducial points (nasion, left and right preauricular points, inion, and Cz) and to map the contour of the participant’s head. These markers were then employed to refine the registration between the template MRI and the EEG-MEG sensors using an iterative alignment algorithm. For source modeling, we used the ‘Overlapping Spheres’-method to create a cortical surface head model for the MEG data, and a Boundary Element Model (BEM) with Open-MEEG for the EEG data [HML99; Gra+10]. We estimated the noise covariance matrix from a one-minute silence recording made immediately before the audio book session for each participant. For source reconstruction, we applied minimum norm imaging (using the sLORETA measure) with constrained dipole orientations [Pas+02]. We first performed source reconstructions on the averaged ERPs and ERFs for each participant, followed by a grand-average of these reconstructions across the exploration and validation datasets.

### 2.5 Statistical Tests

To systematically assess differences in ERFs and ERPs, we performed a cluster-based paired t-test on all 29 subjects (two-tailed, 5,000 randomizations) employing the FieldTrip toolbox [MO07] within the Brainstorm environment. This analysis used the averaged ERF/ERP data from each participant, spanning the -1.0 to 2.0 s window around word onset across all sensors and channels.

In sensor space (method *timelockstatistics*), we compared the 29 averaged ERFs/ERPs for nouns and verbs to identify significant differences. In source space (method *sourcestatistics*), we reconstructed word-type-specific ERFs for each participant and compared these averages with those derived from randomly selected time points matched for trial number. We then computed the mean activity within each of the 600 regions defined in the Schaefer2018 MNI parcellation [Sch+18; Yeo+11]. For visualization, we generated a global source map illustrating the significant p-values (p *<* 0.05) across all parcellations, allowing clear identification of brain-regions with contributions to the significant differences.

We further examined potential predictive coding effects in the data by performing a paired permutation test across specific brain areas, comparing averaged noun and verb ERPs/ERFs to random time points in sensor space [Pan+05]. This two-tailed Wilcoxon signed-rank test (5,000 randomizations) was controlled for false detections across signal, time, and frequency dimensions (FDR correction). In the MEG data, we concentrated on left frontal channels (A229, A212, A178, A154, A126, A230, A213, A179, A155, A127, A177, A153, A125). In the EEG, analyses are focused on the left and right temporal channels (P11, TP7, TP9, M1, P12, TP8, TP10, M2).

### 2.6 Analysis of LLM LLaMa 3.2

We used the pre-trained multilingual large language model Llama-3.2-1B (version release date: September 25, 2024, [AIM24]) as a computational platform, extracting hidden representations from both its embedding layer and each of its 16 transformer blocks using PyTorch [Pas+19]. Llama incorporates a rotary position encoding (RoPE) mechanism to capture token order, while its initially random embedding matrix was refined during pre-training, potentially encoding both semantic and syntactic cues [AIM24; Tou+23b; Tou+23a]. The model has a vocabulary size of over 128 k tokens, can process sequences of up to 128 k tokens, and was trained on approximately 9 trillion tokens for next-token prediction [AIM24].

To gain further insights into the mechanisms underlying predictive coding, we conducted two additional experiments utilizing the LLM Llama to explore the relationship between its internal representations and the neural signals observed in this study. This approach aims to shed light on whether - and how - artificial models replicate the brain’s ability to anticipate specific linguistic features, thereby refining our understanding of how both biological and computational systems manage predictive processing in language. Accordingly, our primary research question is whether prediction-related neural activity is more closely associated with the semantic context of the narrative — referred to here as “semantic prediction” — or whether it is more influenced by the knowledge of the preceding word’s grammatical category, defined here as “syntactic prediction.”

In the “semantic prediction”-experiment with Llama, we used the pre-trained Llama model and fed in chunks of the audio book with increasing length step-by step to the LLM, limiting the context to the 200 preceding words to reduce computational load. Each time, we used PyTorch’s softmax function [Pas+19] to compute the probability of the actual next word. If a word was split into multiple subtokens, we averaged their probabilities. This approach allowed us to evaluate the model’s ability to predict individual words (see Fig. 7 J) based on the preceding text segments. Thus, we calculated a prediction probability for each individual word in the text. For instance, given the input “The Hyades lie in the center of the …”, Llama predicted the next word in the text - “circle” - with a probability of 0.17 (see Fig. 7 J). We repeated this for all words in the audiobook and averaged the prediction probabilities based on the predicted word classes.

In the “syntactic prediction”-experiment, we fed the entire segment of the audio book as presented to participants into Llama at once and read out the hidden-representations of the transformer blocks (Fig. 7 K). We utilized hooks to capture the embeddings of tokens at several stages: after the embedding layer of Llama (layer 0) and following each decoder layer (layers 1-16). Without additional fine-tuning, we extracted the hidden representations for each word. In cases where words were split into subtokens, only the first embedding was stored as an approximation for the entire word. We then assessed how well these hidden representations clustered according to the grammatical class of the subsequent word and quantified this separability using the Generative Discrimination Value (GDV) [Sch+21b]. To further analyze these embeddings, we trained a probe classifier to predict the grammatical class of the next word based on the current word’s embedding, i.e. we used the word class of the next word as label for the embedding input (Fig. 7 K). We employed a simple linear model architecture for probing comprising a linear layer (input dimension = 2048, hidden dimension = 512), a ReLU activation function, and a final linear layer (hidden dimension = 512, output dimension = 4). Given the highly imbalanced dataset, we sub-sampled all word classes to match the smallest category (325 for adjectives) and partitioned the data into training (70%), validation (15%), and test (15%) sets. This allowed us to assess the extent to which information about word classes is already encoded in the hidden representations, incorporating more complex non-linear relationships.

## 3 Results

### 3.1 ERFs and ERPs in Sensor Space

To ensure reproducibility, we split the recorded data into an exploration set (19 subjects) and a validation set (10 subjects) before analysis. This allowed us to verify that observed temporal and spatial signals reflected word-type effects rather than artifacts or noise. Striking similarities in sensor-level MEG and EEG average responses between both sets confirmed the stability and robustness of these language-related signals. Furthermore, direct comparisons of ERFs and ERPs showed no significant differences between the exploration and validation datasets (see Fig. 2 for ERFs and Fig. 3 for ERPs), further confirming the reliability of our measures across participant groups.

**Figure 2:**
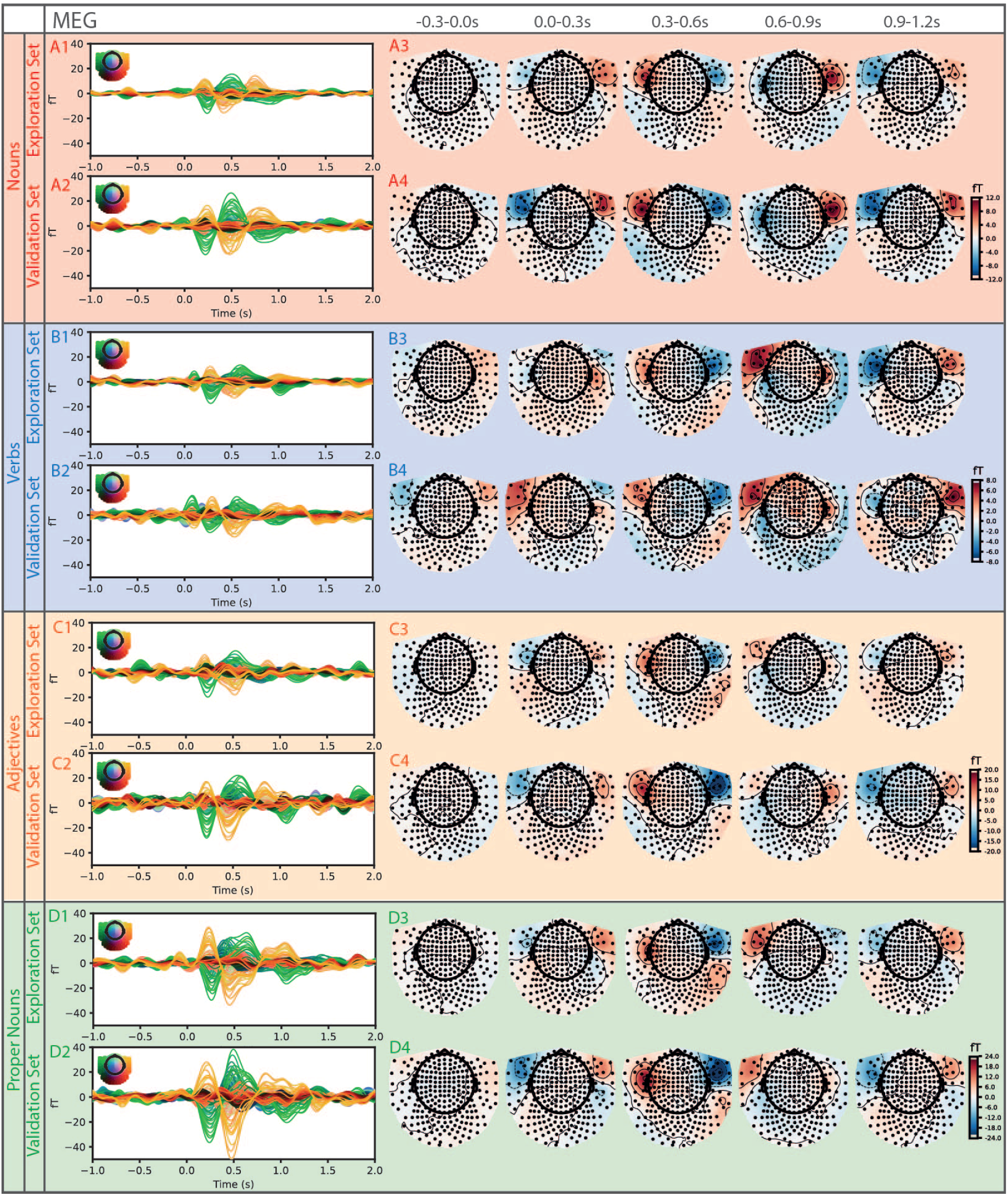
Grand average ERFs (MEG) in sensor space for different word types split into exploration (subject 1-19) and validation (subject 20-29) sets. A1: Average nouns ERFs in exploration set with average topographic maps in five time frames (-0.3-0.0 s, 0.0-0.3 s, 0.3-0.6 s, 0.6-0.9 s, 0.9-1.2 s) in A3. A2: Average ERFs induced by nouns in validation set with corresponding average topographic maps in A4. B,C,D 1-4: Same analysis as for nouns, for verbs (B), adjectives (C), and proper nouns (D)

**Figure 3:**
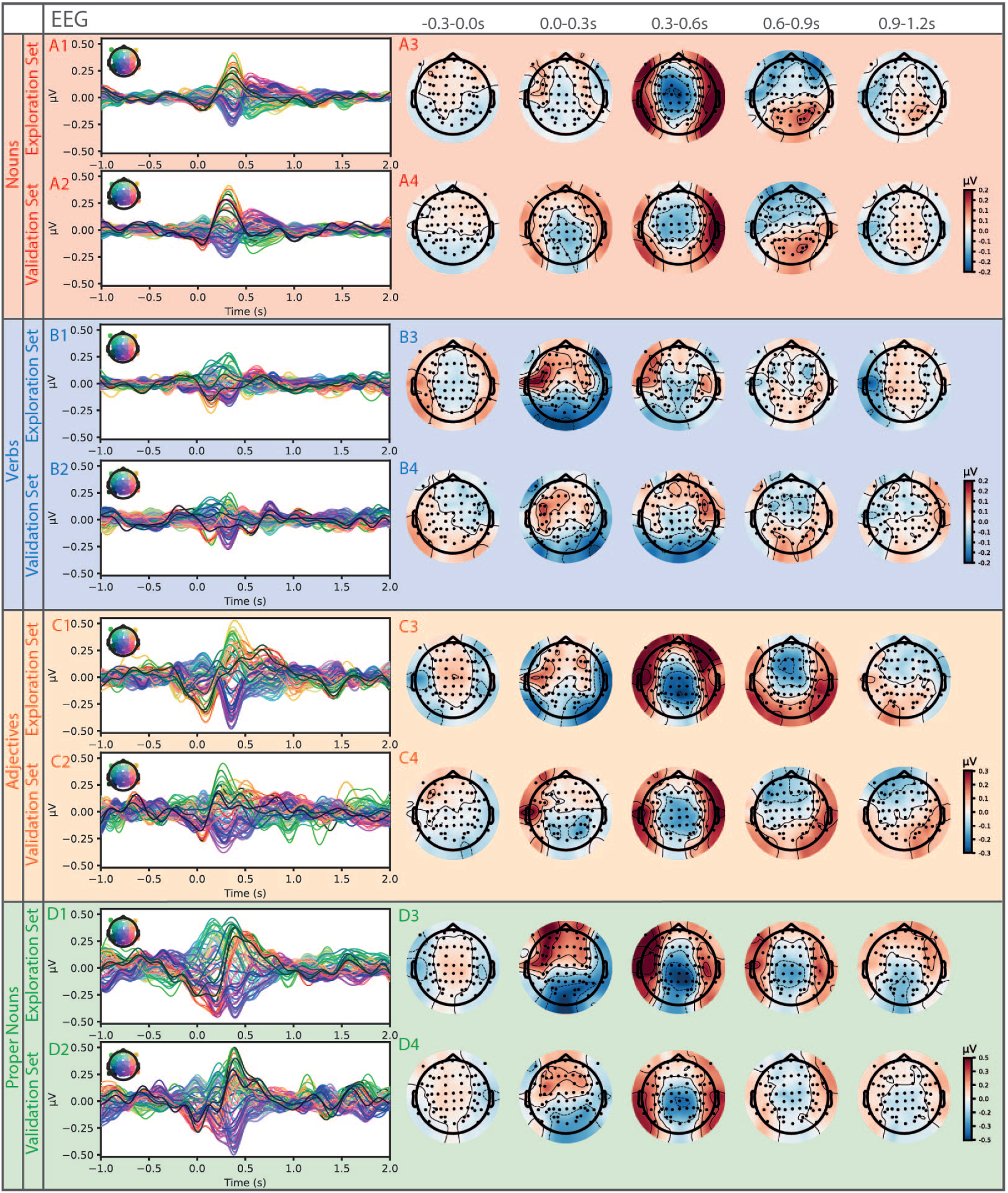
Grand average ERPs (EEG) in sensor space for different word types split into exploration (subject 1-19) and validation (subject 20-29) sets. A1: Average nouns ERPs in exploration set with average topographic maps in five time frames (-0.3-0.0 s, 0.0-0.3 s, 0.3-0.6 s, 0.6-0.9 s, 0.9-1.2 s) in A3. A2: Average ERPs induced by nouns in validation set with corresponding average topographic maps in A4. B,C,D 1-4: Same analysis as for nouns, for verbs (B), adjectives (C), and proper nouns (D)

The corresponding topographic maps generated from the MEG data, which emphasize the spatial distribution of the signal (see Fig. 2 A,B,C,D, 3-4), also show strong concordance between the exploration and validation data sets for all four word types. In the time interval between 0.3 and 0.6 s after word onset, a robust signal pattern emerges across word types, characterized by left hemisphere positivity and right hemisphere negativity in frontal regions. In contrast, the subsequent 0.6-0.9 s window shows more word-type-specific differences and corresponds to the typical latency of the P600 wave, often associated with syntactic processing [Kaa+00]. Within this interval, proper nouns evoke the strongest amplitudes, followed by adjectives, nouns, and verbs.

**Figure 4:**
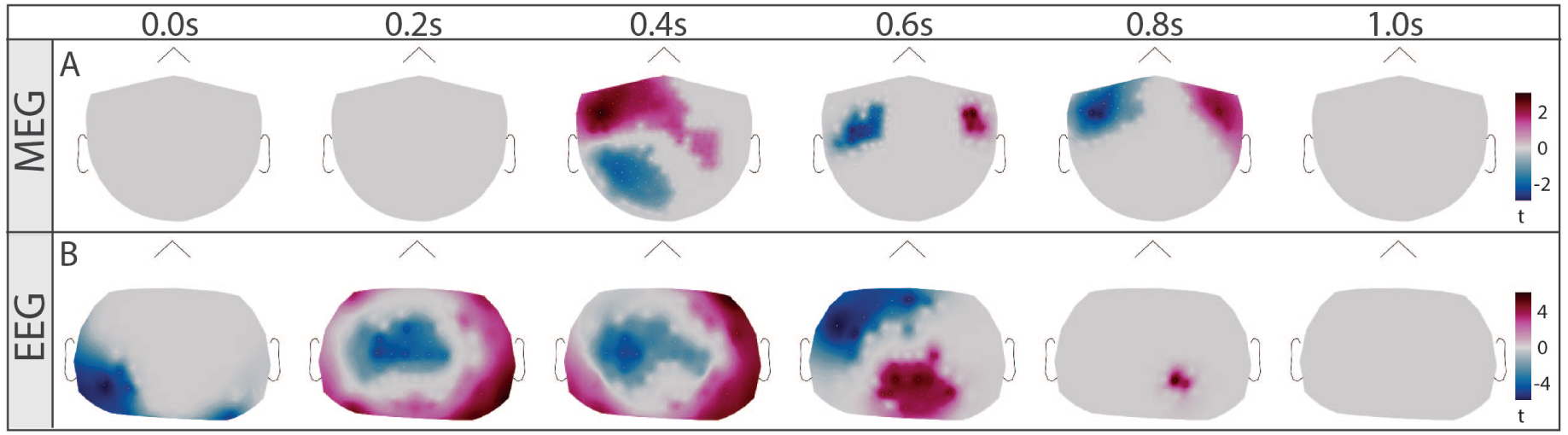
Significant differences of spatial distribution of ERFs and ERPs (topographic maps) between nouns and verbs based on cluster-based paired t-test ([MO07]) (time interval: 0 - 1 s, shown time points: 0.0 s, 0.2 s, 0.4 s, 0.6 s, 0.8 s, 1 s). Significant p-values of clusters in MEG: red cluster: p = 0.0008, blue cluster: p = 0.0012. In EEG: red and blue cluster p = 0.0004.

In analogy to MEG topographic maps, the EEG data also show strong similarity between the exploration and validation data sets across all word types (see Fig. 3 A,B,C,D, 3-4). The N400, widely associated with semantic processing, clearly emerges in these data [LPP08]. Again, proper nouns and adjectives show the largest N400 amplitudes, while verbs exhibit the weakest. All word types, except verbs, show a comparable bilateral response in the time interval between 0.3-0.6 s after word onset, which then diverges in the following window. In contrast to the MEG findings, the EEG data signal amplitude is lower in the time interval between 0.6 and 0.9 s.

To estimate the signal-to-noise ratio of the actual signal, we compared the ERFs and ERPs with pseudo-ERFs/ERPs calculated from random time points (see Suppl. Fig. 1 for random time points, for word types ERFs/ERPs against baseline see Suppl. Fig. 2 and 3). As the audio book contained far fewer adjectives (#325) and proper nouns (#413) than verbs (#936) and nouns (#1376), the resulting lower number of trials produced a poorer signal-to-noise ratio - particularly noticeable in the EEG data. As a result, we focused on nouns and verbs in the subsequent analyses. We used cluster statistics to compare the spatio-temporal patterns associated with verb and noun processing and identified significant distinctions in both sensor-level EEG and MEG data (see Fig. 4) (for comparisons of other word types see Suppl. Fig. 4 (MEG) and 5 (EEG)). In the EEG data, clusters were observable immediately at word onset (0.0 s), providing a first indication for predictive coding (Fig. 4 B) [SBF21]. In contrast, the MEG data revealed significant differences from 0.4 s post-onset (Fig. 4 A). Between latencies of 0.4 s and 0.8 s we found significant distinctions between verbs and nouns in MEG as well as in the EEG data.

### 3.2 Source Space Analysis

To map the neural activity to the exact brain regions we performed source space analysis. First, we visually verified the source-reconstructed ERFs and ERPs for both the exploration and validation data sets for nouns and verbs to ensure consistency and reliability in our spatial analyses (see Fig. 5 (A-D) for MEG and (E-H) for EEG, RMS amplitudes in intervals 0-0.3 s, 0.3-0.6 s, and 0.6-0.9 s). However, the relative sparseness of the EEG recordings (64 channels) compared to MEG recordings (248 sensors) limits the precision of the source reconstructions, leading to differences between exploration and validation datasets (cf. e.g., Fig. 5 E and F at 0 s-0.3 s). Consequently, we focused our source space analyses on the MEG data only, taking advantage of its superior spatial resolution to precisely localize brain activity associated with specific grammatical structures. MEG source space reconstructions showed high reproducibility between the exploration and validation datasets, identifying significant neural sources in the temporal lobes with a pronounced left lateralization during the 0-0.3 s interval (Fig. 5 0-0.3 s). In addition, sustained activity emerged in both hemispheres from 0.3 to 0.9 s after word onset, indicating a dynamic bilateral engagement of brain regions involved in grammatical processing (Fig. 5 0.3-0.9 s). To further identify the brain regions involved in processing different word types, we used cluster-based statistical analyses to contrast the source-space signals elicited by nouns and verbs with those derived from averaging responses at random time points.

**Figure 5:**
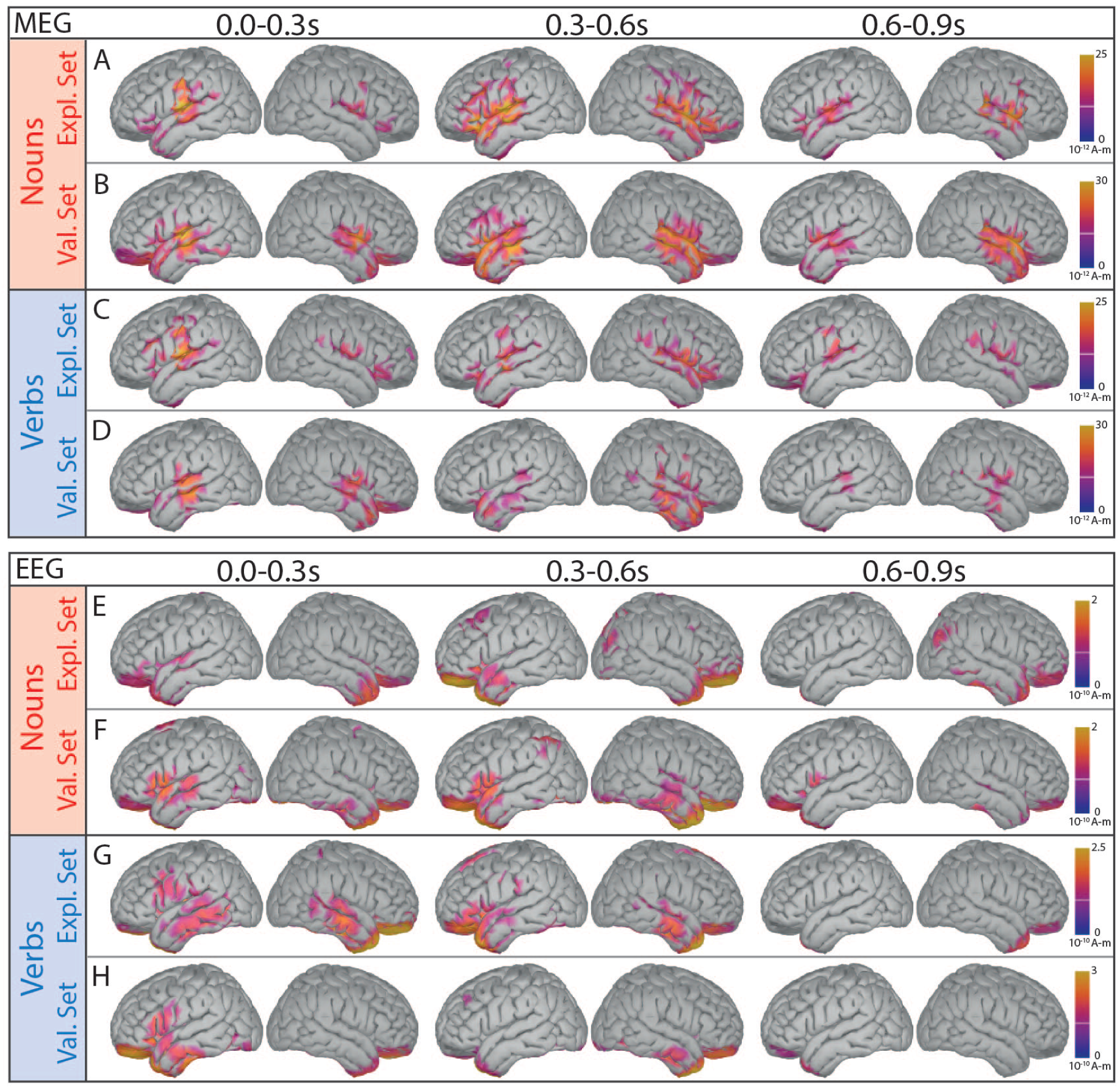
Source reconstructed ERFs/ERPs in three time intervals (RMS amplitudes in intervals: 0.0-0.3 s, 0.3-0.6 s, 0.6-0.9 s). Grand average ERFs (MEG) of nouns for exploration (A, subjects 1-19) and validation data set (B, subjects 20-29). C-D: Source reconstructions for verbs. E-H: Source reconstruction for ERPs (EEG) for nouns (E-F) and verbs (G-H).

The activity is considerably stronger for nouns compared to verbs from 0.3 s until 0.9 s (see Fig. 5 A vs. C). Most strikingly, besides frontal and temporal activations, significant activation was observed in the precentral and postcentral gyri during noun processing for latencies higher than 0.6 s (see Fig. 6 A). Conversely, verbs predominantly activated the left and right subcentral gyri (see Fig. 6 B).

**Figure 6:**
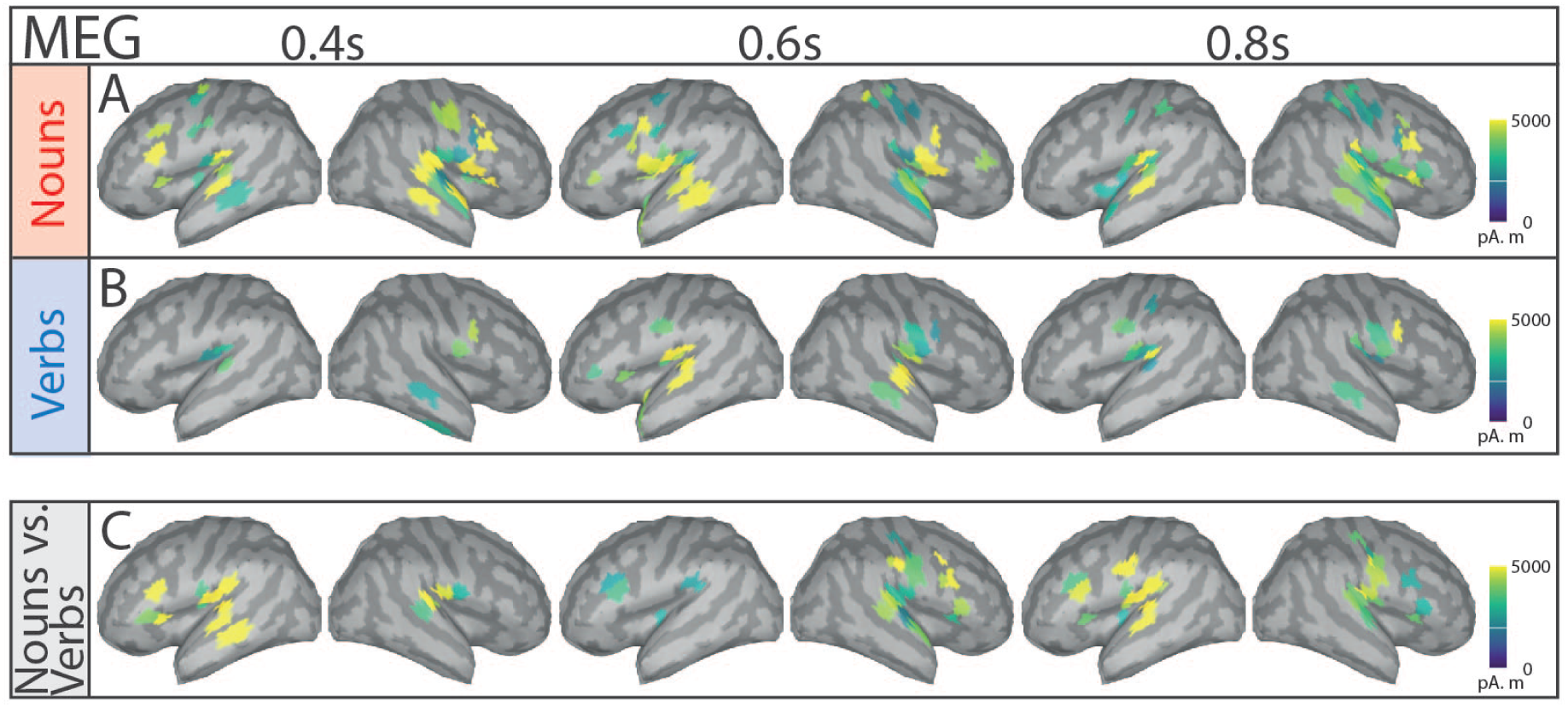
Significant differences of source space activity between nouns and baseline, verbs and baseline, and nouns and verbs based on cluster-based paired t-test ([MO07]) (shown time points: 0.4 s, 0.6 s, 0.8 s). Shown amplitudes is simulated MEG activity in pAm using the resulting t-values of significant regions (p*<*0.05). A: Noun-ERFs vs. random-time point ERFs (significant difference from baseline), B: Verb-ERFs vs. random time point ERFs and C) Noun-ERFs vs. verb-ERFs.

Thus, nouns and verbs engage distinct neural mechanisms, with nouns eliciting higher signal amplitudes in the precentral and postcentral cortex regions (somatosensory, motor, and pre-motor cortex). This aligns with the embodied cognition framework, suggesting that language and sensory systems are interconnected [PHH01; Bar08]. Nouns may carry more meaning on average than verbs, though verb types vary in semantic richness, contrasting with Maess et al.’s claim that verbs hold more semantic information [Mae+16]. Additionally, verbs help predict upcoming nouns [TKH21], warranting further study to advance understanding of semantic grounding and embodied cognition.

Disentangling semantic and syntactic processing in the brain using continuous speech stimuli presents a difficult challenge. To gain a deeper understanding of these mechanisms, it is crucial to further investigate brain signals associated with prediction.

### 3.3 Predictive Coding

To systematically investigate predictive coding in the brain, we next focused on the temporal dynamics of verb and noun processing. Accordingly, we returned to sensor space and performed paired permutation tests on the MEG data (Fig. 7 A, C, E, G), evaluating averaged signals from left frontal channels for each word type to determine whether significant activity emerged around 0 s. A corresponding analysis in the EEG data (see Fig. 7 B, D, F, H) targeted combined left and right temporal channels. In the EEG, nouns, adjectives, and proper nouns elicited a significant negative peak beginning before 0 s, whereas verbs showed no evidence of early predictive activity. The MEG data also revealed a significant peak around 0 s in left frontal channels for nouns, but not for the other word types, suggesting early predictive mechanisms specifically associated with this word class. To gain further insights into the mechanisms underlying predictive coding, we conducted two additional experiments utilizing the LLM Llama to investigate the correspondence between its hidden representations and the neural signals observed in this study.

**Figure 7:**
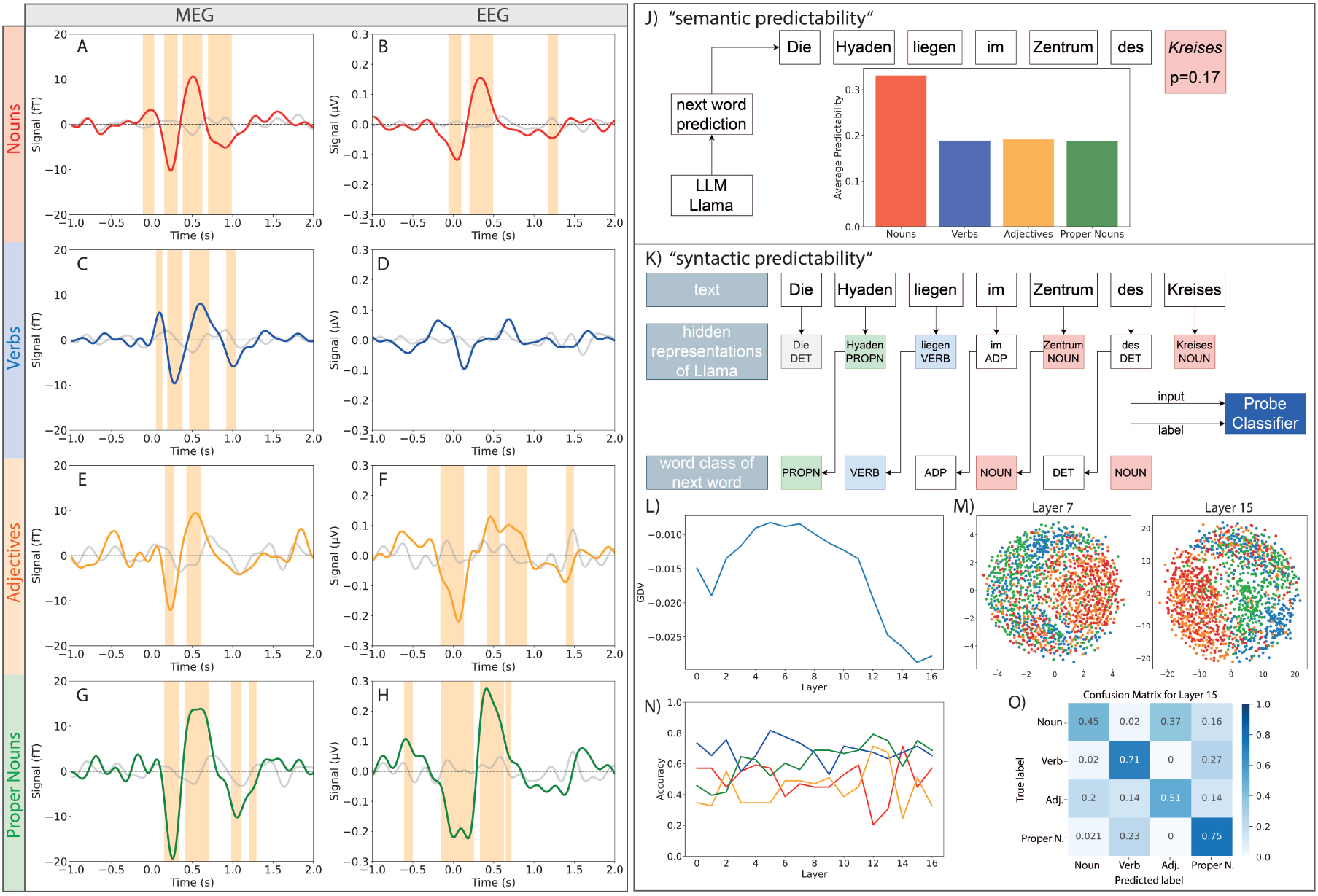
Predictive coding in ERPs and ERFs and in the LLM Llama. A: Permutation-test (FDR corrected, nouns vs. baseline, red) ([Pan+05]) on ERFs averaged across left frontal channels (’A229’, ‘A212’, ‘A178’, ‘A154’, ‘A126’, ‘A230’, ‘A213’, ‘A179’, ‘A155’, ‘A127’, ‘A177’, ‘A153’, ‘A125’). C, E and F Verbs(blue)/Adjectives(orange)/Proper Nouns(green) against baseline for the same sensors as in A. B: Permutation-test (FDR corrected, nouns vs. baseline, red) on ERPs using average signal across temporal channels (’P12’, ‘TP7’, ‘TP9’, ‘M1’, ‘P11’, ‘TP8’, ‘TP10’, ‘M2’). D, F, H: Verbs (blue)/ adjectives (orange)/ proper nouns (green) vs. baseline for same EEG sensors as B. Significant (p*<*0.05) time intervals marked in orange. J) “semantic predictability”: The probability for each word in the audio book was calculated using next word prediction with LLM Llama. Averaging the probabilities of all nouns, verbs, adjectives and proper nouns demonstrated that nouns are easiest to predict by semantic context. K) “Syntactic predictability” based on the hidden representations of Llama for each word embedding as input data and the word class of the next word as labels for training a linear probe classifiers. L) GDV scores of the representations for each layer show a first increasing and then decreasing curve with minimum at layer 15. MDS plots (M) display embeddings projected on 2 dimension for layer 7 (left) and layer 15 (right). (N) Accuracy of the probe classifier of each word class, and (O) confusion matrix of layer 15. The probe classifier shows preformance above chance level. Verbs and proper nouns are clearly separated while adjectives and nouns show a significant overlap.

As result of the “semantic predictability”-experiment we found that the prediction probability for the next word is highest for the word class “noun”, and considerably lower for the other three word classes (see Fig. 7 J). However, in the EEG data the prediction signal is similarly high for nouns, adjectives, and proper nouns (see Fig. 7 B, F, G). Therefore, the underlying mechanisms in the brain might differ from those in the LLM.

In the second experiment - “syntactic predictability” - we calculated the separability of the hidden representations of Lama according to the word class of the next word (see Fig. 7 K). Analysis of the discrimination value shows that already in the embedding layer the word representations (embeddings) cluster according to the word classes of the subsequent words (see Fig. 7 L). Visualization of 2D projections of the embeddings are shown in Fig. 7 M. These embeddings should not contain much information on the context of the actual story, as the context is first added in the attention-heads of the LLM ([Tou+23b; Che+23]). The pre-training should already have added some information on the syntactic structure of sentences to the embeddings. Therefore, this experiment was developed to test the prediction on syntactic rules what we call the “syntactic prediction”. Note, however, that syntax and semantics are not strictly separable in LLMs and that this distinction used here as an approximation with the purpose to gain more insights in semantic and syntactic processing. We could show that across the transformer blocks the clustering of proper nouns considerably increases (see green dots in Fig. 7 M). Proper nouns are located and used similarly in sentences as nouns in general. However, the fact that proper nouns and nouns cannot be easily separated in the early transformer blocks compared to the later ones, indicates that this separability depends on the semantic content. By using a probe classifier trained on the hidden representations we see that the (prediction-) classification accuracy does not change much across the transformer blocks. However, accuracy consistently increases exclusively for the proper nouns (see Fig. 7 N). This is likely due to the semantic content of the proper nouns. We analyzed the confusion matrix for the different word classes and found that adjectives and nouns are often confused (Fig. 7 O). This can potentially be because nouns and adjectives are used at similar positions in sentences in the german language. Thus, it is e.g., impossible to predict purely from syntactic considerations, how the sentence “We have picked up a …” has to be completed and if the next word is a noun or an adjective. It is possible that the next word is a noun: “We have picked up a signal” or an adjective: “We have picked up a strong signal” (Fig. 7 O). An analysis of the audio book text confirms this, as 80% of all adjectives appear preceding nouns within sentences. Whether these findings are generalizable to other languages remains to be tested.

The fact that the prediction-related amplitude of the ERPs is similar for nouns, proper-nouns and adjectives might indicate that simple syntactic rules are integrated in the temporal regions of the brain. In contrast, ERFs show greater differentiation between proper nouns and nouns, which may suggest that frontal brain regions integrate multi-model information with memory content - possibly indicating involvement in semantic processing.

We acknowledge that this represents only a preliminary step toward understanding syntactic and semantic processing by comparing LLMs and human brain activities. We present initial evidence suggesting that different transformer blocks may functionally correspond to distinct cortical regions, though further exploration is needed.

## 4 Discussion

### 4.1 Summary

In this study, we simultaneously recorded EEG and MEG data from 29 healthy German-speaking participants while they listened to an audio book, thus capturing neural responses to continuous naturalistic speech. After signal filtering and segmentation pre-processing, we computed event-related fields (ERFs) and event-related potentials (ERPs) for four word types - nouns, verbs, adjectives, and proper nouns - and performed both sensor- and source-level analyses to characterize the spatio-temporal dynamics of speech resp. language processing. Our main findings reveal characteristic spatio-temporal activation patterns for different word types, likely reflecting both syntactic and semantic processes. In source space, nouns elicited pronounced activation not only in temporal and frontal areas, but also in the precentral and postcentral gyri, suggesting that they may carry a heavier semantic load [Koe25]. Furthermore, we observed significant pre-onset activity exclusively for nouns and adjectives in both EEG and MEG, whereas no comparable predictive signals emerged for verbs. In the temporal-lobe-ERPs we also found significant pre-onset activity for proper nouns, whereas in the frontal-lobe-ERFs we found no activity before proper noun onset.

In addition, we performed a series of experiments with a LLM (Llama-3.2-3B) and compared the findings with the predictive neuronal data of MEG and EEG measurements. In our first experiment, we evaluated what we call “semantic prediction” of the pre-trained Llama-3.2-3B model by presenting it with progressively longer segments of audio book text to assess its ability to anticipate the subsequent word from the context. Across the dataset, nouns were predicted with a higher probability than other word classes. However, the fact that nouns are predicted far better than other word classes does not fit to what we see in the neuronal data and thus we did further experiments with Llama.

In our “syntactic prediction” experiment row, we fed an extended segment of the audio book into Llama-3.2-3B and extracted hidden representations from all 16 transformer blocks [AIM24]. We then quantified the separability of these representations using the Generalized Discrimination Value (GDV) to find out how much predictory power is stored in the hidden representations of certain words. Our analyses showed that even at the embedding level, the model’s representations began to cluster by grammatical class, with this separation becoming more pronounced in later transformer blocks. In particular, proper nouns and nouns became increasingly distinct, suggesting that syntactic information is progressively integrated with semantic information throughout transformer blocks. As an extension of the syntactic prediction experiment, we trained a linear classifier (probe network, [Bel22]) on Lama’s hidden representations to assess the extent to which information about upcoming word classes was encoded in non-linear relationships, which were already covered by the GDV analysis. These probe experiments revealed that prediction accuracy remained relatively stable across transformer blocks, with improved accuracy for proper nouns, suggesting that syntactic and semantic cues are progressively integrated during language processing in accordance to the results described above. Our findings suggest that the similar prediction-related ERP amplitudes (EEG) for nouns, adjectives and proper nouns in temporal regions indicate the integration of simple syntactic rules in analogy to the embedding layer of Llama. In contrast, the different ERF patterns (MEG) observed in frontal regions - particularly between proper nouns and nouns - suggest that these areas may integrate more complex multi-modal information into memory, likely reflecting semantic processing.

### 4.2 Activity in sensory and motor system as indication for semantic grounding

Our work explores the relationship between semantic grounding, multi-modal integration in language understanding, and our findings in the source space analysis. The activity patterns observed in the precentral and postcentral cortex evoked by nouns but not verbs may result from covert sensorimotor processes rather than overt motor behavior (Fig. 6). In our study, no motor tasks, visible motor cues, or tactile stimulation were provided, eliminating voluntary actions and mirror-related activity as potential sources of the motor cortex response [Che85; DD10; NLP14]. Although participants were instructed to remain still-potentially suppressing movement-related signals-such suppression would likely manifest in a temporally uncorrelated manner [EB17]. Instead, the prominent motor and somatosensory activations detected in the absence of explicit movement tasks could be explained by engagement in motor imagery (MI) [MP21; Sin+24]. Indeed, it has already been shown that the ventral precentral gyrus, related to tongue movement for speech production could be used for brain computer interfaces (BCIs) in a sense that a cursor could be moved just by imagining and mentally verbalizing the action [Sin+24]. Consistent with theories of embodied cognition, previous studies have shown that action-related language processing can evoke activity in motor regions as early as 200-400 ms post-stimulus [Bon+22; Pul+05; PHH01]. These activations may stem from the heavier semantic load nouns carry relative to verbs, a distinction that should also shape the predictability of these word classes. Consequently, examining word-class-specific predictability could help disentangle the extent to which neural responses reflect semantic grounding in contrast to purely syntactic processes.

### 4.3 Prediction of different word classes in the brain and LLMs

From the perspective of the Bayesian brain and especially of the free energy principles, the cognitive system continuously refines its internal model of language to minimize prediction error and surprise when anticipating the next word [WES24]. Thus, the brain continuously generates predictions about incoming sensory inputs, including the next word during language comprehension [SK24; FK09; SBF21]. As Grisoni and coworkers have shown, before presenting a critical word, context-induced semantic predictions are reflected by a semantic readiness potential (SRP), which is the neural correlate of that prediction. However, this SRP is only present before words with high-constraint context [GMP17; GBP24]. To determine whether the predictive signal in our data is consistent with semantic readiness potentials observed in studies with less naturalistic stimuli, we compared the hidden representations of the LLM Llama with the observed neural patterns. Our analyses show that the predictive signal in the brain seems to consist of two components: a syntactic readiness, which can be observed in temporal regions, and a semantic readiness potential, mainly located in frontal areas, consistently with previous reports [GMP17]. These convergent findings are consistent with the Bayesian framework of the brain, suggesting that both neural and computational systems continually update their internal models by integrating prior expectations with incoming information. Taken together, this evidence highlights the presence of distinct but complementary predictive mechanisms underlying semantic and syntactic processing in language comprehension.

### 4.4 LLMs as a model for the brain

The question remains whether LLMs can serve as a valid model for understanding the human brain. Large language models are constructed as layered stacks of transformer blocks that operate via self-attention rather than explicit recurrent connections [Tou+23b; Tou+23a; Fer+25], yet their repetitive structure - characterized by self-similarity and fractal organization being a universal principle in biological structures [Gui+16; La +18; Zhe+20]- may serve as an analogue to the recurrent transmission of signals through the arcuate fasciculus. In the human brain, the arcuate fasciculus facilitates dynamic bidirectional communication between Broca’s and Wernicke’s areas, a pathway long proposed as the neural substrate for a universal innate grammar [Fri+17; ZT21]. Although LLMs do not replicate the full complexity of biological recurrence, the iterative processing achieved by stacking transformer blocks appears to approximate the brain’s integration of syntactic and semantic cues. These observations suggest that the self-similar structure of the LLM may provide valuable insights into the neural strategies underlying predictive coding and integrative processing during language comprehension.

### 4.5 Conclusive Remarks

Our study demonstrates that predictive coding in language processing operates through both syntactic and semantic anticipation, as reflected in distinct pre-word onset activity captured by combined MEG and EEG recordings. To overcome the limitations of conventional averaging and low signal-to-noise ratios, future research in cognitive computational neuroscience should incorporate advanced single-trial analyses and deep learning techniques to improve word class classification and reveal underlying grammatical structures (see, e.g., [Mis+24]). In addition, our results suggest that LLMs provide a computational framework that approximates human predictive coding, with their stacked transformer blocks potentially mirroring the recurrent interactions between Broca’s and Wernicke’s areas via the arcuate fasciculus. The self-similar organization of these transformer architectures may reflect universal hierarchical principles of cognitive processing, shedding new light on the computational basis of linguistic prediction. The integration of neurophysiological data with advanced AI models provides a promising framework to disentangle the interplay between syntactic structure and semantic context in predictive coding. The fusion of generative AI and neural data has the potential to refine cognitive computational neuroscience (CCN, [KD18]) and provide deeper insights into the hierarchical organization of language processing in biological and artificial systems. Ultimately, bridging neuroscience, AI and linguistic theory may not only reveal the cognitive mechanisms that govern human language, but also drive the development of artificial intelligence - bringing it closer to the way the human brain anticipates and processes language [Sch+23a; Has+17].

## 5 Acknowledgments

This work was funded by the Deutsche Forschungsgemeinschaft (DFG, German Research Foundation): grants TZ 100/2-1 (project number 510395418), KR 5148/2-1 (project number 436456810), KR 5148/3-1 (project number 510395418), KR 5148/5-1 (project number 542747151), and GRK 2839 (project number 468527017) to PK, and grant SCHI 1482/3-1 (project number 451810794) to AS. Furthermore, the research leading to these results has received funding from the European Research Council (ERC) under the European Union’s Horizon 2020 research and innovation programmme (ERC Grant No. 810316 to AM).

## 6 Author Contributions

AS and PK developed the study protocol. AS, PK, SR, KT, AM supervised the study. NK and MK performed the measurements. NK developed the evaluation programs. All authors discussed the results. AS, PK, NK drafted the first version of the manuscript. AM and TK reviewed the first draft of the manuscript. All authors accept the final version of the manuscript.

## 7 Supplementary Material

Additional figures are presented in this section, offering deeper insights into the EEG and MEG analyses. Figure 8 illustrates an example of average baseline activity, generated using randomly selected time points with an equivalent number of trials as those for nouns. The signal amplitudes for ERFs (left) and ERPs (right) are much lower compared to those for word-class-specific ERFs and ERPs in Figure 3 and 2.

**Figure 8:**
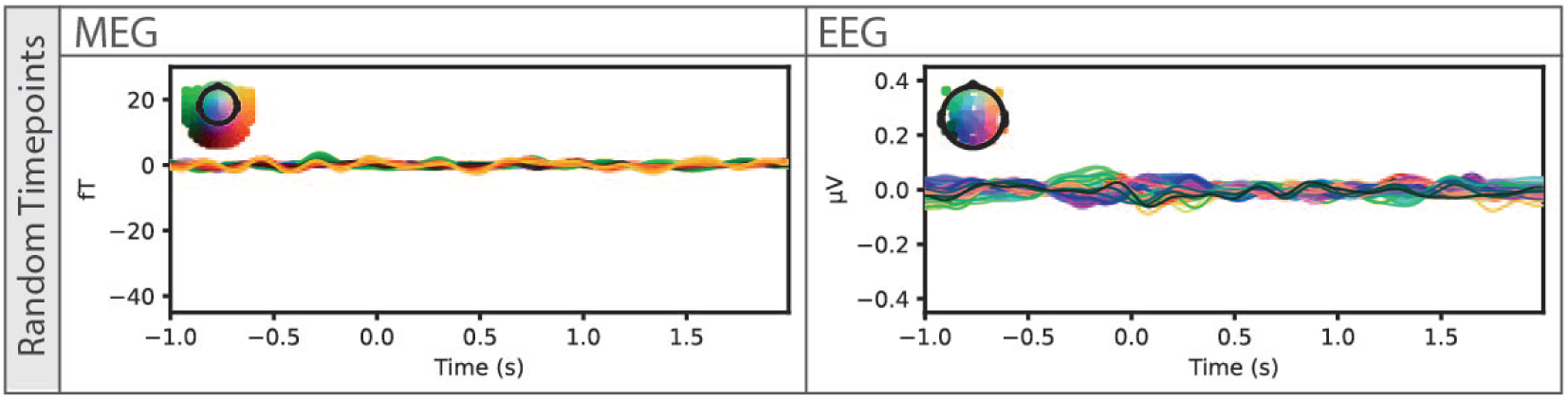
Grand average ERFs (left) and ERPs (right) of brain activity at randomly chosen time points. Example shows the signals for the same amount of trials as for the nouns.

Figures 9 (ERFs) and 10 (ERPs) depict significant clusters identified through a cluster-based paired t-test, comparing word-class ERFs and ERPs against the baseline. The corresponding six cluster tests between each word class pair (nouns vs. verbs, nouns vs. adjectives, nouns vs. proper nouns, verbs vs. adjectives, verbs vs. proper nouns, and proper nouns vs. adjectives) are shown in Figure 11 for MEG and Figure 12 for EEG. All tests indicate significant differences between word types, with the exception of the ERFs comparison between verbs and adjectives.

**Figure 9:**
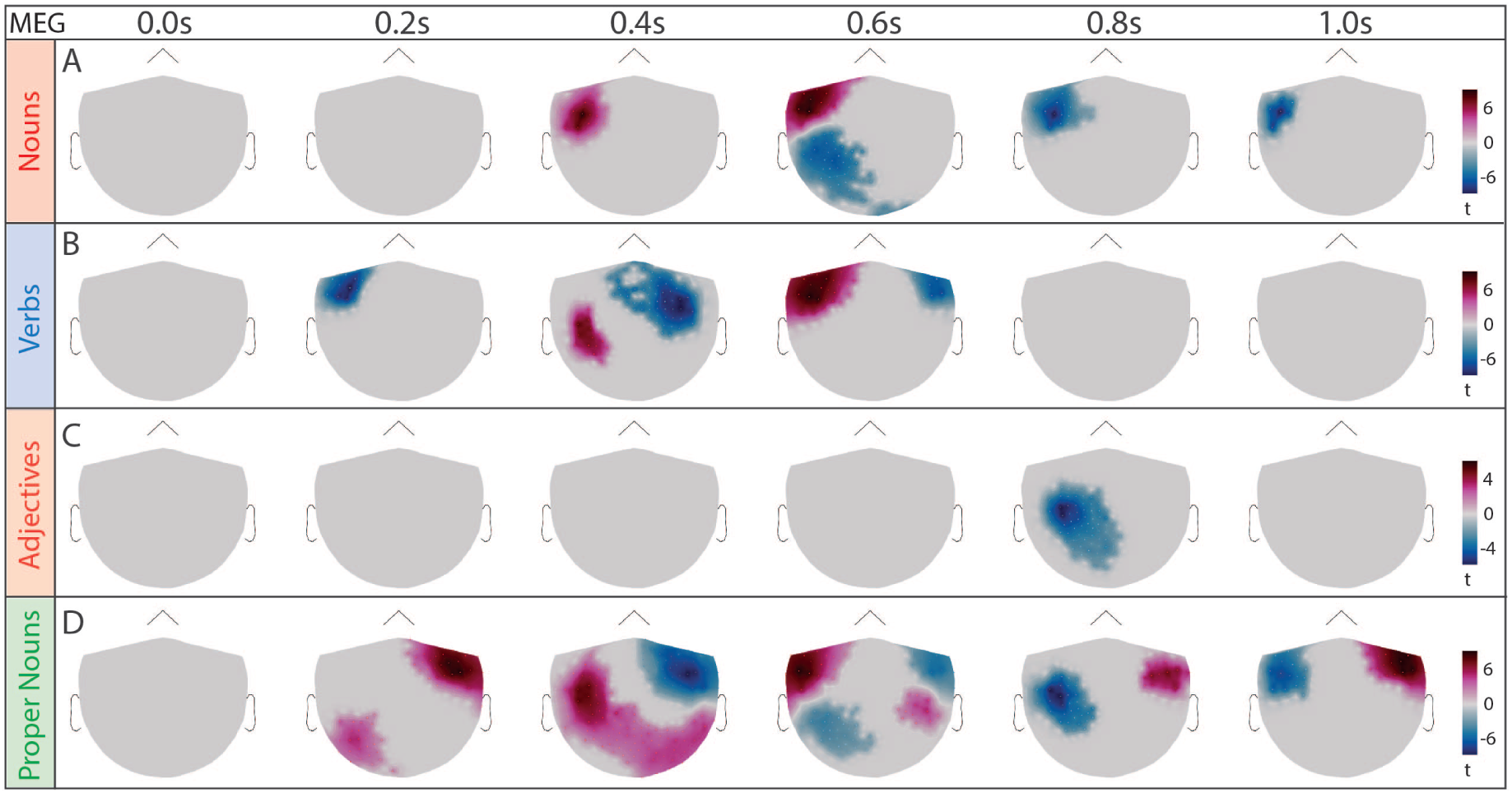
Significant differences of spatial distribution of ERFs (topomaps) between word classes (nouns, verbs, adjectives, proper nouns) and baseline based on cluster-based paired t-test (time interval: 0-1s, shown time points: 0.0 s, 0.2 s, 0.4 s, 0.6 s, 0.8 s, 1 s). A) nouns: red: p=0.018, blue: p=0.00004. B) verbs: red: p=0.0328, blue: p=0.0008. C) adjectives: red: p=0.0488, D) proper nouns: red: p=0.0004, blue right hemisphere: p=0.0120, blue left hemisphere: p=0.0020.

**Figure 10:**
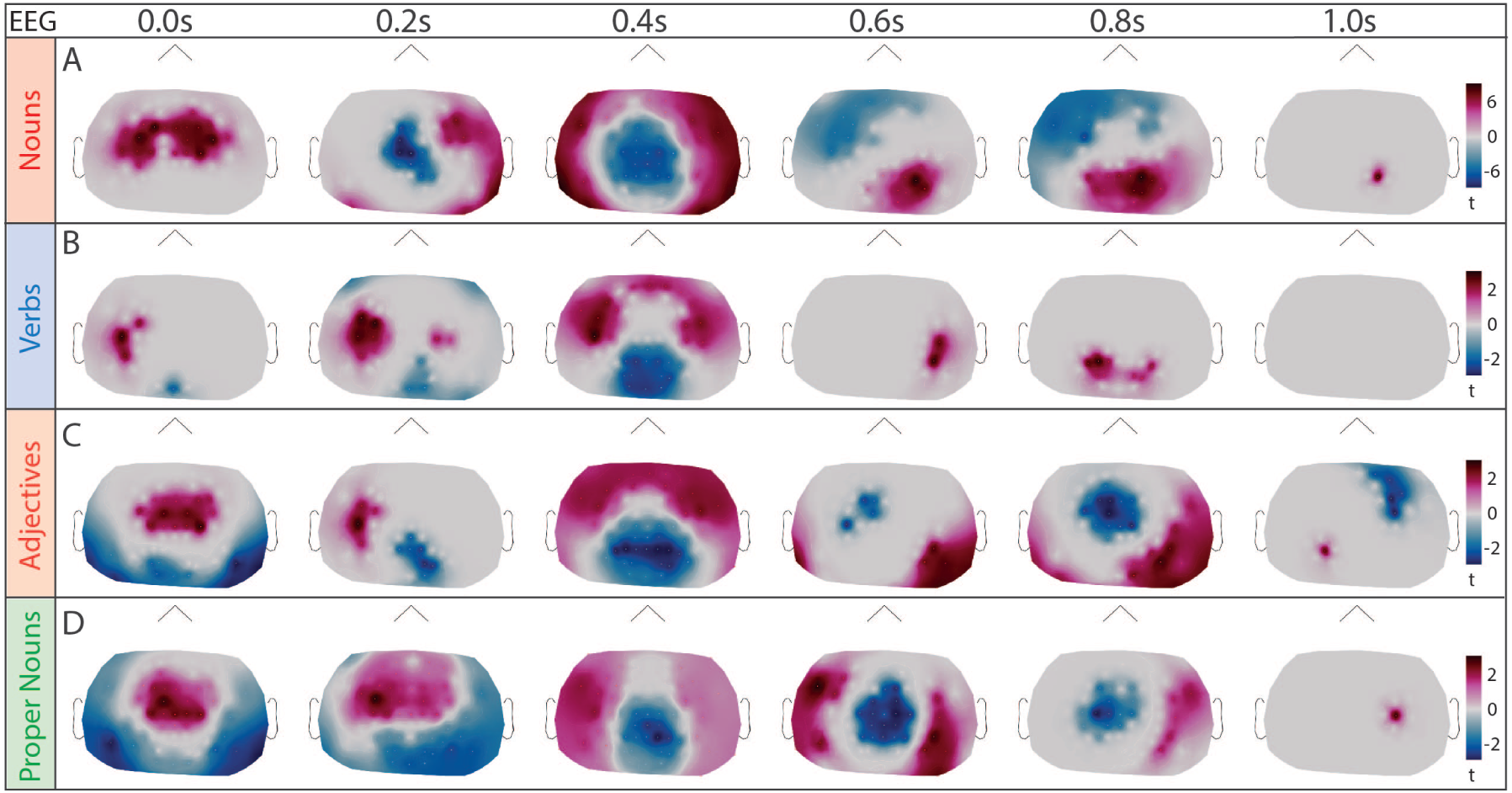
Significant differences of spatial distribution of ERPs (topomaps) between word classes (nouns, verbs, adjectives, proper nouns) and baseline based on cluster-based paired t-test (time interval: 0-1s, shown time points: 0.0 s, 0.2 s, 0.4 s, 0.6 s, 0.8 s, 1 s). A) nouns: red: p=0.0004, blue: p=0.0004. B) verbs: red: p=0.0004, blue: p=0.0028. C) adjectives: red: p=0.0004, blue: p=0.0004. D) proper nouns: red: p=0.0004, blue: p=0.0004.

**Figure 11:**
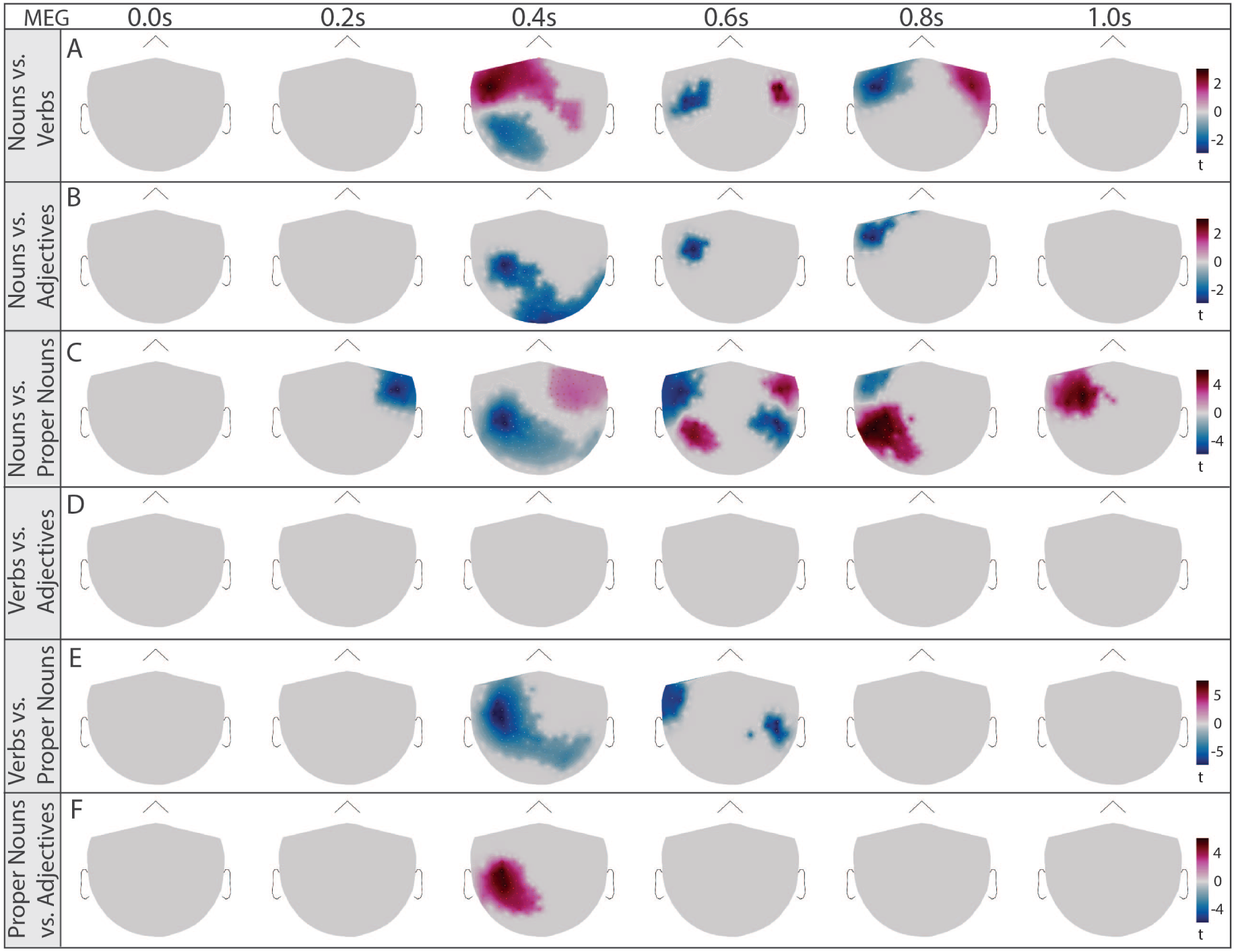
Significant differences of spatial distribution of ERFs (topomaps) between all word classes (nouns, verbs, adjectives, proper nouns) based on cluster-based paired t-test (time interval: 0-1s, shown time points: 0, 0.2, 0.4, 0.6, 0.8, 1s). Significant clusters found in five investigations. A) nouns vs. verbs: blue: p=0.0012, red: p=0.0008, B) nouns vs. adjectives: blue: p=0.0080, C) nouns vs. proper nouns: red left hemisphere: p=0.0004, red right hemisphere: p=0.0296, blue: p=0.0004, E) verbs vs. proper nouns: blue: p=0.0004, F) proper nouns vs. adjectives: red: p=0.0210.

**Figure 12:**
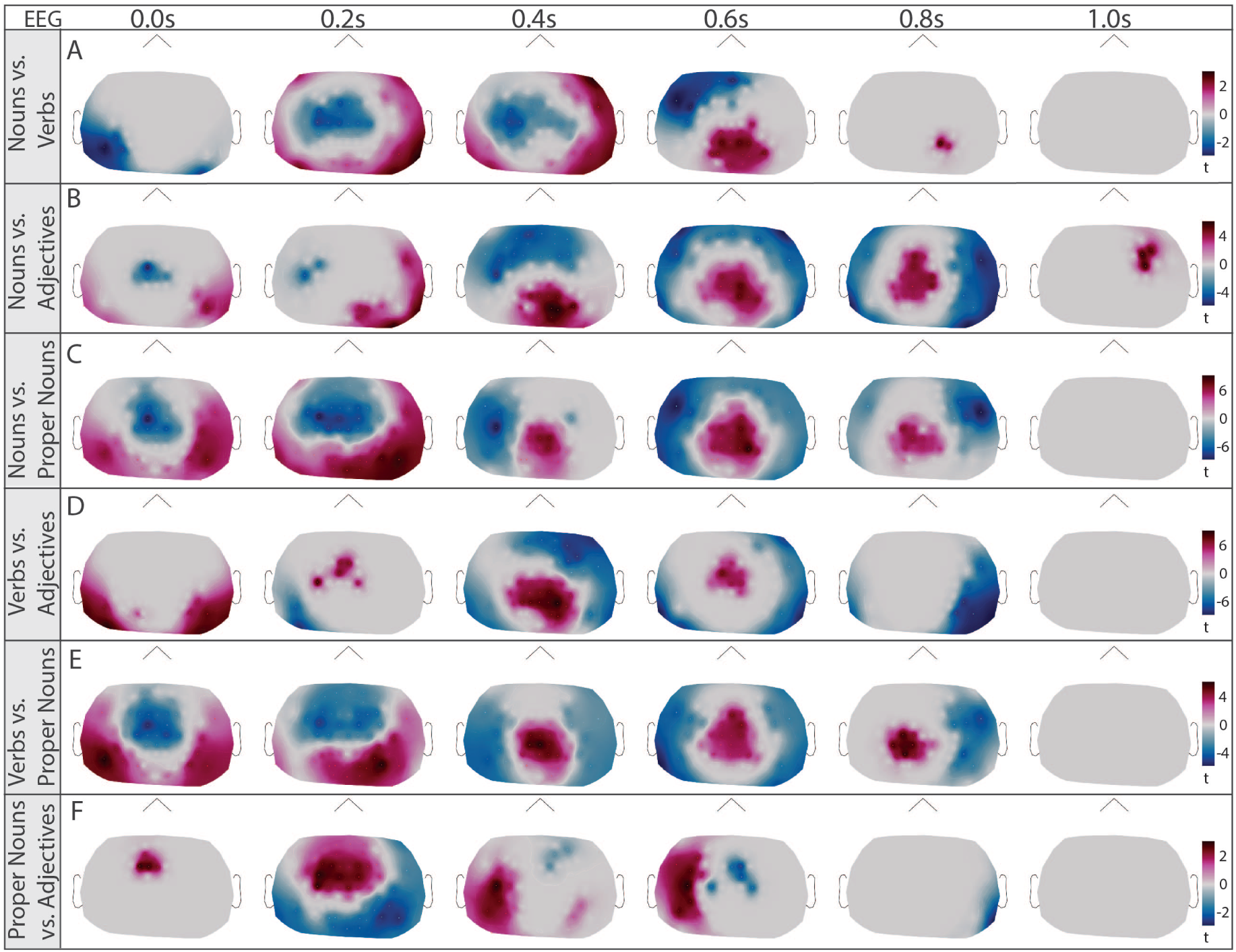
Significant differences of spatial distribution of ERPs (topomaps) between all word classes (nouns, verbs, adjectives, proper nouns) based on cluster-based paired t-test (time interval: 0-1s, shown time points: 0, 0.2, 0.4, 0.6, 0.8, 1s). A) nouns vs. verbs: blue: p=0.0004, red: p=0.0004. B) nouns vs. adjectives: blue: p=0.0004, red: p=0.0004. C) nouns vs. proper nouns: red: p=0.0004, blue: p=0.0004. D) verbs vs. adjectives: blue: p=0.0004, red: p=0.0004, E) verbs vs. proper nouns: red: p=0.0004, blue: p=0.0004. F) proper nouns vs. adjectives: red: p=0.0024, blue: p=0.0004.

The poorer source reconstruction for EEG data can be demonstrated by the lack of differences in source-space activations for nouns and verbs in Figure 13 compared to Figure 6.

**Figure 13:**
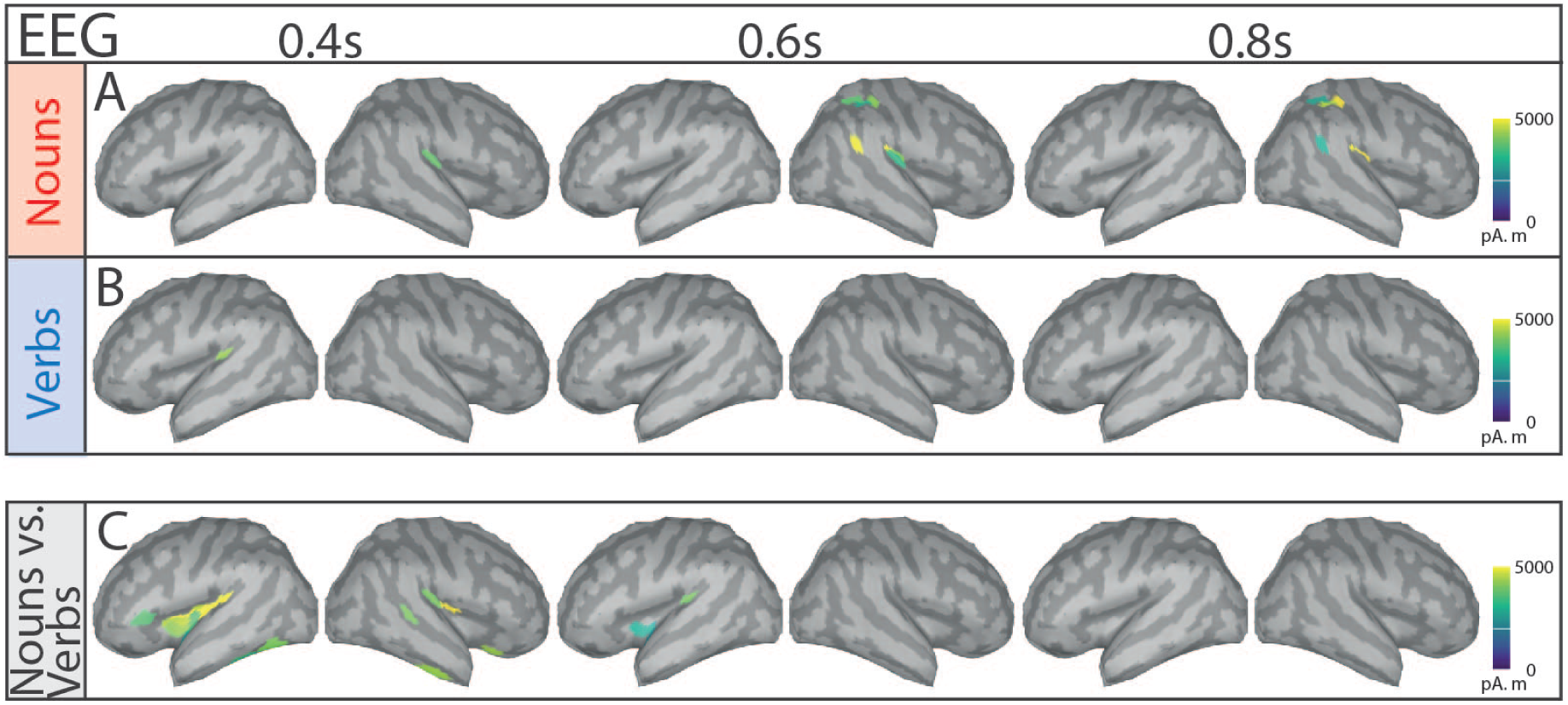
Significant differences of source space activity between nouns and baseline, verbs and baseline, and nouns and verbs based on cluster-based paired t-test (shown time points: 0.4s, 0.6s, 0.8s). Shown amplitudes is simulated MEG activity in pAm using the resulting t-values of significant regions (p<0.05). A: Noun-ERPs vs. random-time point ERPs (significant difference from baseline), B: Verb-ERPs vs. random time point ERPs and C) Noun-ERPs vs. verb-ERPs.

